# A multiparametric, quantitative and high-throughput assay to quantify the influence of target size on phagocytic uptake

**DOI:** 10.1101/482547

**Authors:** L. Montel, L. Pinon, J. Fattaccioli

## Abstract

Phagocytosis by macrophages represents a fundamental process essential for both immunity and tissue homeostasis. It consists in the uptake of pathogenic or cellular targets larger than 0.5μm. For the biggest particles, the phagocytic process involves a massive reorganization of membrane and actin cytoskeleton as well as an important intracellular deformation, all in a matter of minutes. The study of the role of the size of objects in their phagocytosis has lead to contradictory results in the last decades.

We designed a method using confocal microscopy, automated image analysis and databases for fast quantitative analysis of phagocytosis assays. It yields comprehensive data on the cells and targets geometric and fluorescence intensity parameters, automatically discriminates internalized from external targets, and stores the relationship between a cell and the targets it has engulfed. We used two types of targets, solid polystyrene beads and liquid lipid droplets, to investigate the influence of size on the phagocytic uptake of macrophages. The method made it possible, not only to perform phagocytic assays with functionalized droplets and beads of different sizes, but to use polydisperse particles to further our understanding of the role of size in phagocytosis.

The use of monodisperse and polydisperse objects shows that while smaller monodisperse objects are internalized in greater numbers, objects of different sizes presented simultaneously are internalized without preferred size. Throughout results, the total surface engulfed by the cell appeared to be the main factor limiting the uptake of particles, regardless of their nature or size. A meta-analysis of the literature reveals that this dependence in surface is consistently conserved throughout cell types, targets’ nature or activated receptors.

## INTRODUCTION

Phagocytosis by macrophages represents a fundamental process essential for both immunity and tissue homeostasis. It consists in the uptake of pathogenic or cellular targets larger than 0.5μm. For the biggest particles, the phagocytic process involves a massive reorganization of membrane and actin cytoskeleton as well as an important intracellular deformation, all in a matter of minutes [1]. Influence of biological [2–6], chemical [7–9] and physical [10–13] parameters of the targets on their phagocytosis have been evaluated and compared to unravel the mechanisms of internalization. The main parameter used to assess the efficiency of phagocytosis is the phagocytic index, a tool created by Major Leishman [14], that quantifies the phagocytic process by measuring the average number of internalized objects per phagocyte. Initially, the measurements were performed by counting both cells and targets under a microscope. Since then, methods were devised to obtain the phagocytic index directly from a population of phagocytes, such as spectrophotometry [15–17] or cytometry [18,19], and more recently the counting of cells and targets on microscopy images began to be automatized [20–25]. Phagocytic index, despite its simplicity, is a quantitation averaged over a whole cell population. It is hence not able to capture the full details of the process at the individual level of the cells.

The influence of the size of the targets on the phagocytic efficiency has been studied for more than fifty years [15–19,26–29] for diameters ranging from a hundred nanometers to tens of micrometers. However, experimental results performed with monodisperse particles and published in the literature are contradictory. In some cases, an optimal size for particle uptake seems to emerge [18,27], whereas in others [15,19,28], the number of internalized particles is higher for smaller targets as compared to larger ones in a totally monotonous manner. In some cases, the phagocytic index varies by one order of magnitude for apparently similar experimental conditions reported in two different articles from the literature [16,18,19,27]. Using polydisperse targets could provide new insights about the question of size. However, the phagocytic index, as an averaged quantitation, is inadequate for assessing the phagocytosis of polydisperse droplets. An algorithm that registers the relationships between the targets, and their size, and the cells that have internalized them, is required to evaluate the role of size in the competitive phagocytosis of polydisperse targets.

Herein we have designed a semi-automatic quantitation procedure for image analysis based on confocal microscopy pictures, yielding comprehensive results on the shape, size and fluorescence intensity for both cells and targets. With the aim to answer complex biological questions, results are stored in a relational database that keeps track of all the relationships between cells and targets, and records morphological or biochemical features for each object of the experiment. The method allows relating each cell with its internalized targets in a bidirectional way, hence recording all the available information of a particular experimental condition. Since the segmentation of particles does not require staining, the assessment of the phagocytosis of unaltered particles is possible. Aschematic view of the process is schematized in **Figure 1**.

**Figure 1:**
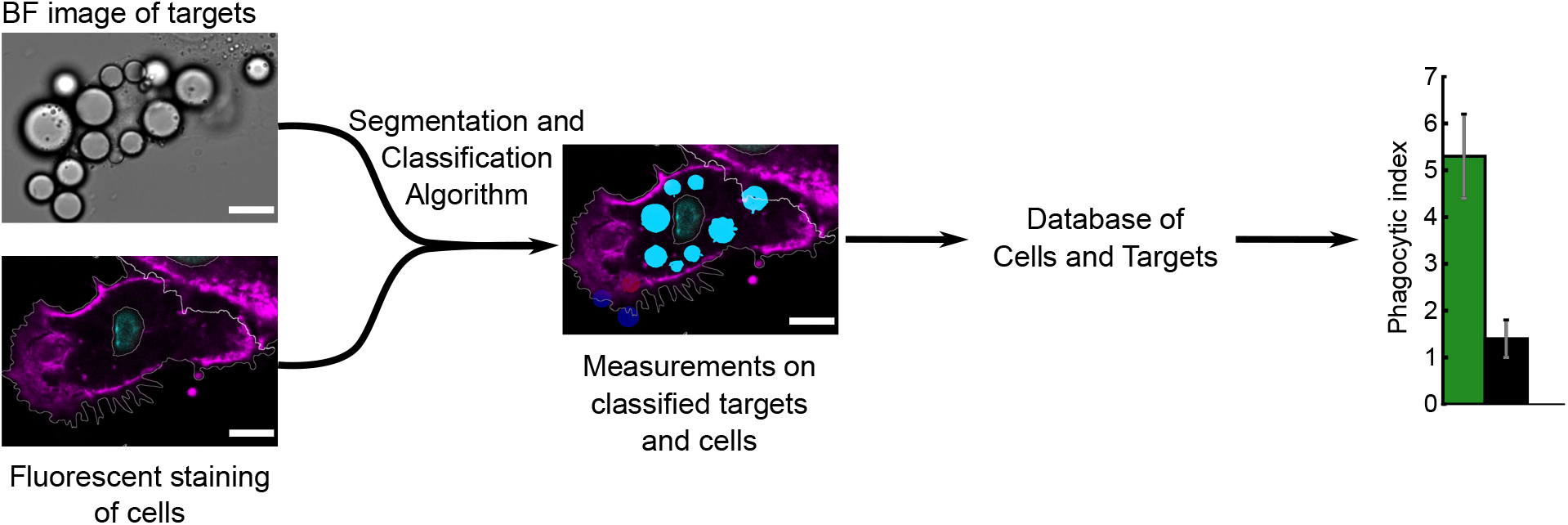
Schematic representation of the data obtained by the automatic detection of phagocytosis. The targets and the cells are detected on the images, classified and measured. The results of the measurements and classification are stored into a relational database where the measurement of the cells, measurement of the targets and the relationship between the targets and the cell they are in. Complex data can be extracted and plotted from this database.

We performed experiments with both monodisperse and polydisperse targets, with and without opsonization, and confronted them with results from the literature about the role of size in phagocytosis. Using the insight provided by the behavior of macrophages towards polydisperse targets, we conclude that the availability of cell membrane surface is the limiting parameter governing phagocytic uptake, regardless of target size. Cells internalize targets indiscriminately until they reach a total internalized surface that differs between non-specific and FcγR-mediated phagocytosis.

## MATERIALS AND METHODS

### Materials

Soybean oil (CAS No. 8001-22-7, Pluronic F68 (CAS No. 9003-11-6), Tween 20 (CAS No. 9005-64-5), sodium alginate (CAS No. 9005-38-3), dimethyl sulfoxide (DMSO) were purchased from Sigma-Aldrich (St Quentin Fallavier, France). Lipiodol (CAS No. 8002-46-8) was a kind gift of the company Guerbet (Villepinte, France). DSPE-PEG-biotin (1,2-distearoyl-sn-glycero-3-phospho ethanolamine-N-[biotinyl(polyethylene glycol)-2000]) were purchased from Avanti Polar Lipids. Alexa Fluor 488-conjugated mouse anti-biotin IgGs (Ref. 200-542-211) were purchased from Jackson Immunoresearch. Polybead^®^ Microspheres (Ref 17134-15, 07312-5 and 17136-5) were purchased from PolySciences (Hirschberg,Germany)

### Emulsion Fabrication

We first disperse by manually stirring 15 g of soybean oil in an aqueous phase containing 2.5 g of a surfactant (Poloxamer F-68, initial proportion of 30 %w/w) and 2.5 g of a thickening agent (sodium alginate, initial proportion of 4 %w/w). This crude, polydisperse emulsion is further sheared and rendered quasi-monodisperse in a Couette cell apparatus under a controlled shear rate, following the method developed by Mason *et al*. [30]. The value of the shear rate determines the size distribution of droplets. Before decantation, the emulsion is diluted in order to have a proportion of 1% w/w of Poloxamer F-68 and 5% w/w of oil. After one night of decantation, the oil phase is get, diluted with a solution of Poloxamer F-68 with an initial proportion of 1% w/w. After several decantation steps to remove very small droplets, the emulsion (final proportion of 50% w/w of oil) is stored at 12°C in a Peltier-cooled cabinet. Size distributions of the emulsion samples are reported in **Figure S1**.

### Emulsion functionalization

The extended version of the functionalization method has already been published [31] and is summarized shortly hereafter. The desired quantity of droplets is washed three times with a solution of Tween20 at CMC (0.06g/l) in Phosphate Buffer Saline. The quantity of DSPE-PEG-biotin required is calculated to be 100 times the number of molecules necessary to saturate the total surface of droplets, based on a minimal occupied area of 12.6 nm^2^ per phospholipid [32]. That amount of DSPE-PEG-biotin saturates the surface of droplets [31] as to prevent the formation of clusters at the surface when the droplets touch a cell. DSPE-PEG-biotin are diluted in DMSO to a total volume of 20μl and mixed with Tween 20 at CMC in PBS and washed droplets for a total volume of 200μl. The suspension is incubated for 90 minutes in agitation at room temperature. Afterwards, the remaining phospholipids and DMSO are diluted at least 10 000 times through 4 washing steps with Tween 20 at CMC in PBS. Droplets are then resuspended with anti-biotin Alexa 488 antibodies (Ref 200-542-211, Jackson ImmunoResearch) in PBS/Tween20 CMC solution, with antibodies in a 1:1 ratio compared to the DSPE-PEG2000-biotin available on the total surface of the droplets and incubated for 45 minutes at room temperature. Remaining antibodies are diluted at least 2000 times through 3 washing steps with PB/Tween20 0.007%. PB/Tween20 is then removed by a final centrifugation step and droplets are resuspended in 40μl/coverslip DMEM 4.5g/L L-glucose without phenol red at 37°C.

### Polystyrene beads functionalization

The desired quantity of beads (2 millions per coverslip) is washed three times with Phosphate Buffer Saline (Ref 14190250, Life Technologies). Beads are then resuspended with FITC antibodies from Human Serum (Ref F9636, Sigma) at 20mg/ml and incubated overnight at room temperature. After centrifugation, supernatant is removed and the beads are washed 4 times in PBS. PBS is then removed by a final centrifugation step and beads are resuspended in 40μl/coverslip DMEM 4.5g/L L-glucose without phenol red (Ref 31053-028, Life Technologies) at 37°C.

### Cell culture

RAW 264.7 murine macrophages (Ref 91062702) were purchased from ECACC (Public Health England, UK). They were cultured in T-80 culture flasks (Ref 734-2131, VWR) with DMEM 4.5g/L L-glucose supplemented with 10% Fetal Calf Serum, 1% Penicillin-Streptomycin and 2mM L-Glutamine (Ref 10313-021, 10500-056, 31053-028 and 25030024 Life Technologies), at 37°C and 5% CO2. 24 hours before experiment, they were detached using TryPLE (Ref 12605-010, Life Technologies) incubated at 37°C for 5 minutes and seeded on 22mm*22mm #1.5 glass coverslips (Ref_631-0125, VWR) in 6-well plates (Ref 734-0054, Corning) at one million cells per well.

### Phagocytosis assay with functionalized droplets or beads

Seeded glass coverslip are mounted on an open chamber formed by two coverslips separated by a 0.1mm thick double-sided tape, and 2 million droplets or beads suspended in 40μl DMEM are injected in the chamber. The droplets and cells are incubated together for 45 minutes at 37°C 5%CO2, then the chamber is rinsed with 160μl PBS and cells are fixed in 4% paraformaldehyde solution.

### Immunostaining

Fixed cells are stained using Hoechst 33258 4μg/ml (Life Technologies, ref H H1399) and phalloidin Atto 550 20μM (Sigma Aldrich, ref 19083) in PBS for 30 minutes.

### Microscopy

Samples were imaged with a confocal Leica TCS SP8, using 2 Hybrid Detectors, and a 40X/1.30NA oil immersion objective. They were illuminated sequentially with four lasers at 405nm, 488nm, 552nm. Hoechst 33258 and Atto550 were observed simultaneously on each of the two detectors, Alexa 488 subsequently. A prism separated the light and the LASX software optimized the window of wavelength gathered for each fluorophore: 410 to 510 nm for Hoechst 33258, 493 to 568nm for Alexa 488, 557 to 789 nm for Atto 550.

### Automated Image analysis

Image analysis was performed with CellProfiler 2 [33] (www.cellprofiler.org) using a custom-built pipeline. All obtained data were stored in a SQL database (SQLite3). Further data visualization was performed with Python3 code using SQLite3 to pass on queries to the database and Matplotlib for plotting. All codes are available upon request.

## RESULTS

We have developed a semi-automatic method for the quantitation of phagocytosis by adherent or semi-adherent phagocytes ingesting various phagocytic probes such as polystyrene beads [34] or lipid droplets [35]. RAW 264.7 murine macrophages [36] were used as the model cell line; and we focused our study on the FcγR-mediated phagocytosis of IgG-coated fluid and solid microparticles, lipid droplets [35] and polystyrene particles [34], respectively.

Bare or IgG-functionalized particulate targets are presented to phagocytic cells for 45 minutes in a cell culture incubator. After rinsing the supernatant, cells are fixed, stained and imaged by confocal scanning microscopy. Both the nucleus and the cytoplasm are stained to perform segmentation of the cells. Here we use Hoechst 33258 [37] and Atto 555-phalloidin [38] as nuclear and cytoskeletal staining agents, respectively. For each sample, corresponding to a glass coverslip, we observe 10 fields of views, each counting at least 300 cells. Consequently, at least 3000 cells per sample are analyzed and quantified for each condition.

The imaging procedure consists in acquiring each color channels at two specific heights inside the sample using a confocal microscope, as schematized **Figure 2A**: (i) at the basal level where the cell is in contact with the glass, referred to as Z_low_; and (ii) at the mid-height level Z_high_, inside the cell, at a vertical coordinate equal to the radius of the targets. Targets, either beads or lipid droplets, appear in the bright field channel as disks with a grey center surrounded by the succession of a white ring and a dark ring at the edge, as visible in **Figure 2B**. These rings result from the difference of refractive index with water [39], which is relatively high in the case of soybean oil (n = 1.47, [40]) and polystyrene (n=1.615, [41]). The cell and its boundaries are easily discernable on the actin cortex imaging at the basal level Z_low_, where the spreading is usually maximal and the intensity more uniform, as shown in **Figure 2C**. By contrast, the area occupied by an internalized target appears dark on the cytoplasmic image recorded at the z=Z_high_ mid-height level, as shown in **Figure 2D**. Dark regions will thus be used afterwards to determine the status of targets regarding their internalization by cells or not.

**Figure 2:**
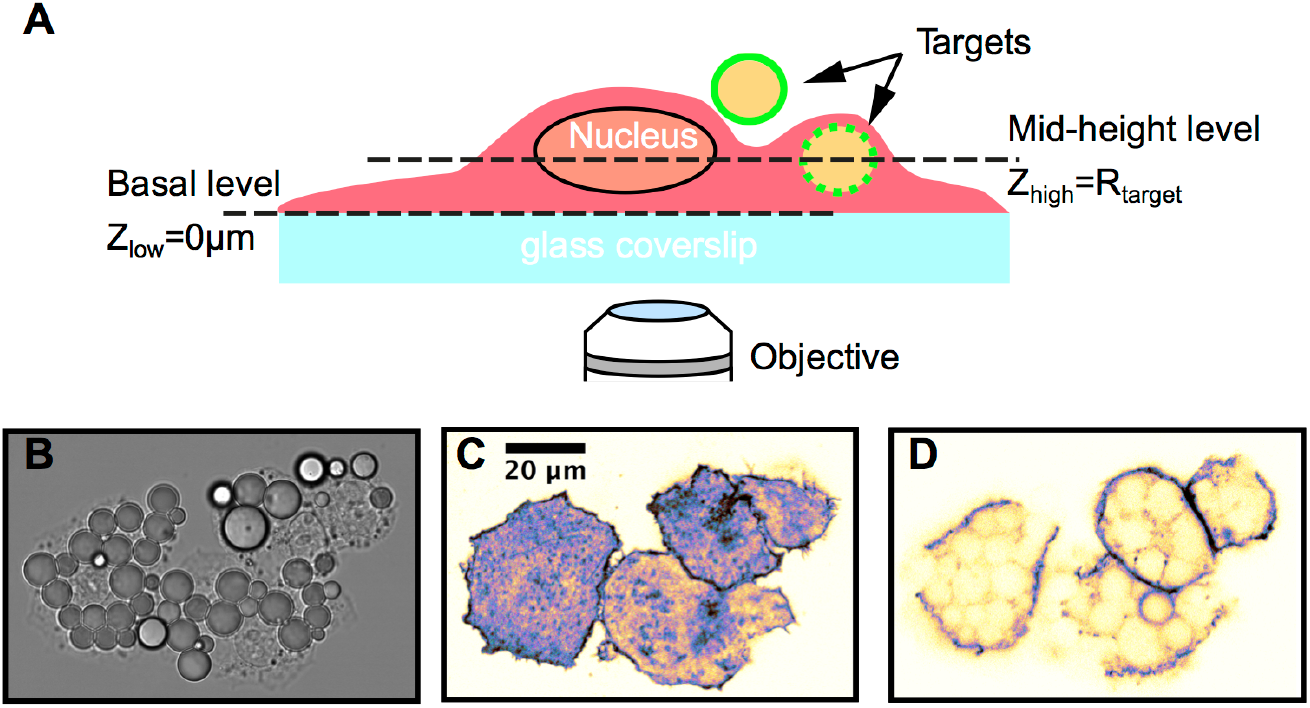
(A) Schematic representation of the two focal planes imaged by confocal microscopy for analysis. (B) Bright field image of macrophages and droplets at Z_high_. (C,D) Confocal image of phalloidin staining of cells at the basal (C) and mid-height (D) levels. The intense basal actin network allows detection of the cell borders, while the holes in the cytoskeletal staining at the places where targets are inside the cytoplasm allow to discriminate internalized targets. Scalebar = 20μm

### Cell segmentation

We use a classical approach to proceed with the cell segmentation, based on localization of the nuclei and on propagation over the basal actin cortex. Nuclei are first segmented from nuclear stained images at the mid-height level Z_high_ (**Figure 3A**) by dividing the image in tiles of one tenth the image size, and applying an intensity threshold obtained by the Otsu method [42] on each tile. Cells are then propagated [43] from the nuclei to their respective cytosol boundaries, over on the actin cortex images recorded at the basal level Z_low_, as illustrated on **Figure 3B**. For individual cells, propagations stop when an abrupt increase on decrease in intensity is detected, in our case a sharp decrease at the border of the cytoplasm. When the propagation of one cell encounters the propagation of a neighbor, the border in determined by a balance between the distance to the nucleus and the intensity gradient at the border. Finally, to get rid of dead cells, cellular debris or potential artifacts, we discard detected objects whose area is under a pre-selected threshold.

**Figure 3:**
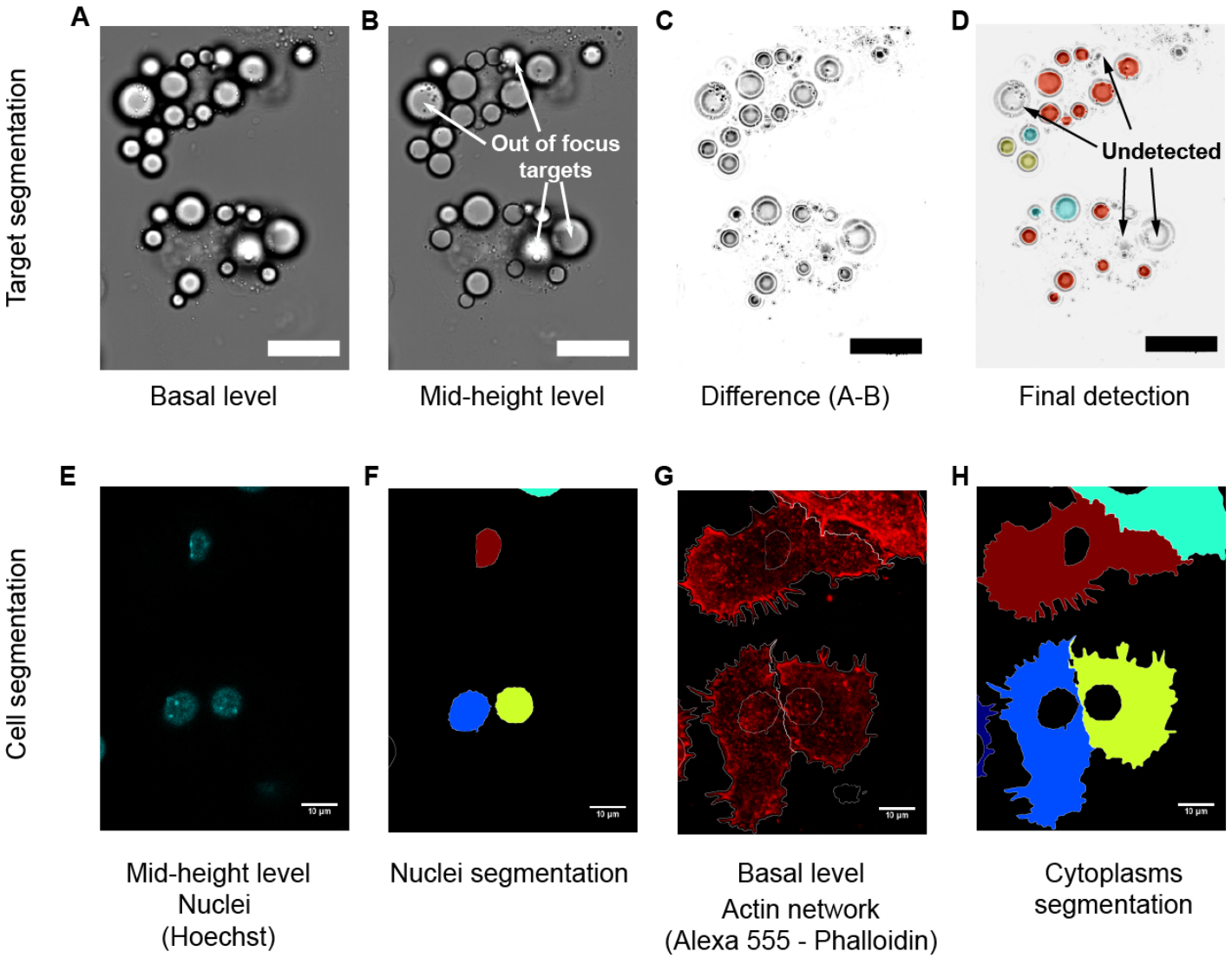
Example of the target segmentation routine (A-D).. The pixelwise difference between the images at two heights (A and B) is calculated (C). On focus targets have different borders on the two images, whereas out of focus targets (arrows) are similar. Thus only on focus targets appear in white on the difference image. The final classification is shown on (D), red being internalized droplets, all other colors external droplets. Steps of cell segmentation (E-H): (E) Nuclei are detected from Hoechst staining at the mid-height Z_high_, (F) each nucleus not touching border is labeled, (G) a propagation algorithm detects the cells border on a phalloidin staining of the actin cytoskeleton at the basal level Z_low_, (H) each cell above the cutoff area is labeled and *Cytoplasm* objects are created by subtracting the nucleus.

The algorithm creates two numerical objects related to the cell cytoplasm, which will be further used for target classification. The first one, called *Cytoplasm*, is created by subtracting the nuclear area from the expanded cell area. The *Cytoplasm* object corresponds to the cytosol as a whole, i.e. including the internalized targets. The second one, called *CytoplasmMinusTargets*, corresponds to the actual cytoplasm of the cells, after having removed the overlapping particles from its envelope. We finally label all *Cytoplasm* and *CytoplasmMinusTargets* objects, record their positions, some geometric features and their relationship to internalized droplets in a database.

### Segmentation of the targets

Targets, either lipid droplets or beads, are segmented from bright field images similar to the one shown in **Figure 3A,B** and **Figure S2**. Two features are detrimental to the correct segmentation and eventual classification of the targets: first, for high phagocytic indexes, particles form clusters from which it is difficult to individualize objects; second, we observe out-of-plane targets at the apex of the cell that may be considered as internalized by classification routine detailed in the next paragraph.

Computing the numerical difference of the bright field images recorded at the basal and mid-height levels solves both issues for oil droplets, as shown on **Figure 3C**. Indeed, we take advantage of the sharp variation of the diffraction rings around the focal plane of the droplets to discriminate droplets in the mid-height plane from droplets above it. **Figure 3C** shows that the borders of the in-plane targets on the two images are very different, whereas the borders of out-of-plane targets are very similar. By computing the difference between the bright field images, we obtain an image where the borders of inplane targets are white, whereas out-of-focus targets appear dark, making the segmentation easy and straightforward. Additionally, the white borders of neighboring droplets are well separated, facilitating the segmentation in crowded regions. To remove small artifacts, we discard acquired targets whose area or form factor is below a pre-selected threshold. The value of both thresholds depends on the size distribution of targets. Conversely, polystyrene microbeads are simply segmented using a pre-defined threshold on the bright field image at the mid-height level Z_high_, thanks to their bright central area. As for cells, we record the position and size of the targets, and store it for further analysis.

### Classification of the targets regarding internalization

For a target to be classified as internalized by a phagocytic cell, we consider and evaluate three independent criteria: (i) the target has to be in the same focal plane than the cell, (ii) the target should be localized within the boundaries of the cytoplasm of a segmented cell, and (iii) most of the area of the target has to occupy a space inside the cytoplasm, meaning that a dark sport is visible in the actin cortex image recorded at the mid-height level. A schematic representation of the classification algorithm is shown in **Figure 4**.

**Figure 4:**
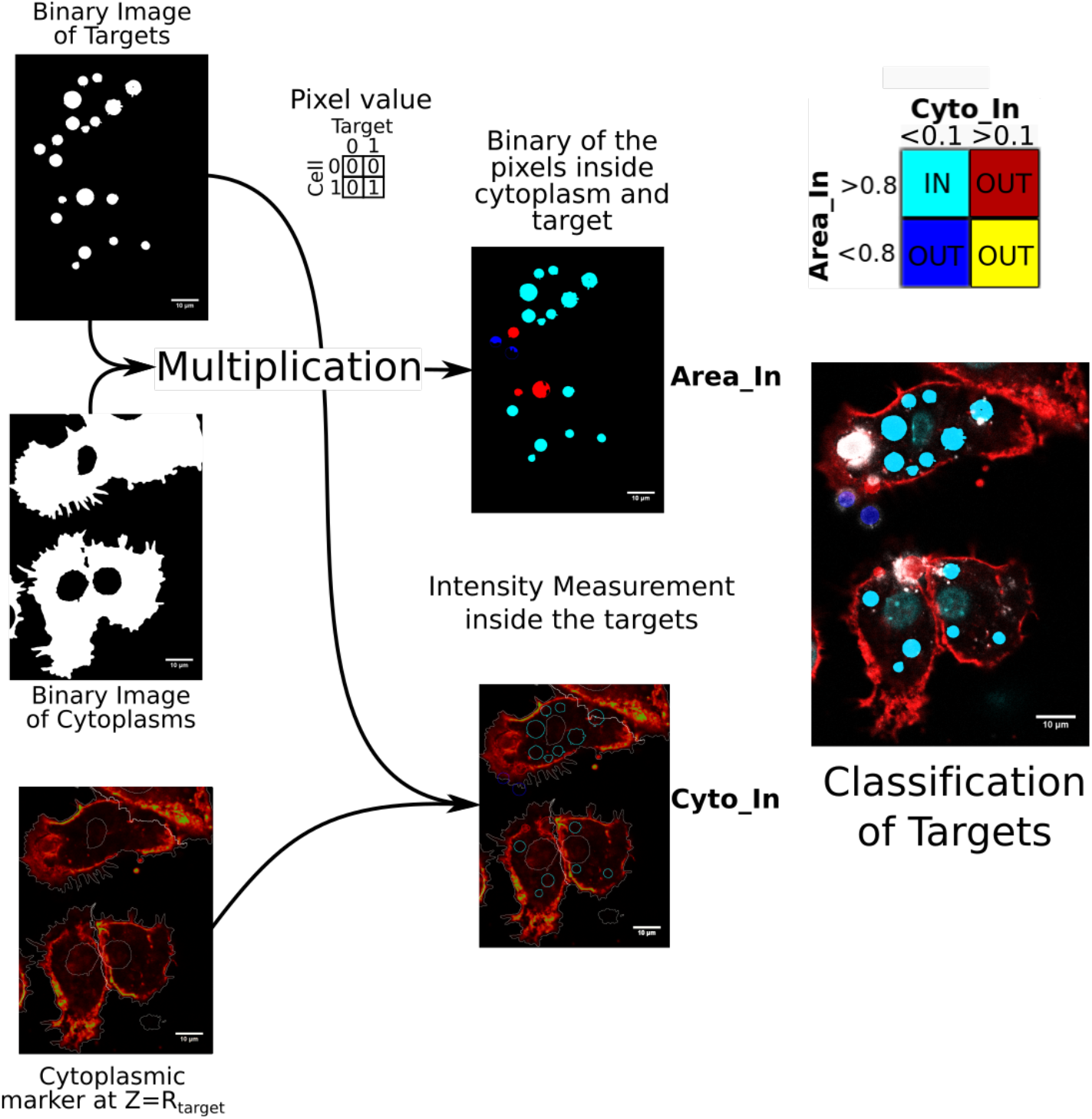
Schematic representation of the classification algorithm that detects internalized targets. Binary images of the segmented targets and cytoplasms are multiplied bitwise so that only the pixels both inside a target and a cytoplasm are 1. The mean value of the pixels of the multiplied image is measured over the whole target, giving a measurement of the percentage of the target area that is inside a cytoplasm. Meanwhile, the value of cytoplasmic staining inside the target is measured. If more than 80% of the target area is in the cytoplasm and the cytoplasmic staining is under a threshold, then the target is deemed internalized (in cyan). Targets displaying too high actin staining (in red) are usually above the cell or being internalized, targets not overlapping the cytoplasm are either outside (in blue) or bordering (in yellow) the cell.

The first criterion is assessed during the target segmentation, where out-of-focus targets are discarded by the segmentation routine. As a consequence, all the detected targets meet this criterion, as those who don’t are discarded prior classification. We assess the second criterion by performing a pixelwise multiplication of the binary image of the cells area and the binary image of the targets area. On the resulting image, the target is considered as internalized following this criterion if more than 80% of its corresponding pixels are located within the border of a single cell. Finally, the third criterion is evaluated on the cytoplasmic image recorded at the mid-height level. The mean intensity of the cytoplasmic staining inside the border of each target is measured after normalization of the fluorescence intensity of the image. Areas characterized by an intensity level higher than 10% of the maximal intensity are classified as non-internalized and discarded.

At the end of the process, internalized targets that fulfill the three above criteria are related to the closest *Cytoplasm* object, i.e. the object that contains them. Their features are measured, and are stored in the database, alongside with the unique identifier of the cell they are in.

### Database structure and population

During the segmentation and classification, morphological features of the cells and targets are acquired, and subsequently stored in a comprehensive SQL database generated by CellProfiler according to our criteria. A diagram of the database is shown on **Figure S3** and details about the fields and categories are summarized in the **Supplementary Information** section. The shape of nuclei, cells and targets is measured from the segmented images. Intensity measurements on all fluorescent channels are performed at the basal and mid-height level for cell cytoplasms. In addition, intensity measurements are performed on targets at the mid-height level.

At the end of this protocol, we obtain a pipeline for phagocytic assays that takes confocal images of the phagocytes and targets as input, and provides a database containing measurements on cells and targets, as well as the relationships between a cell and its internalized targets

### Efficiency of the segmentation routine

To assess the efficiency of the detection and classification algorithm, its results were confronted to a blind manual counting and classification on three randomly chosen images from each of three replicate coverslips. **Table 1** summarizes the results of the algorithm for 3 sizes of beads (3, 6 and 10μm diameter) and 3 sizes of droplets (3, 5.5 and 8μm diameter). Sensitivity, i.e. the part of the targets that are correctly classified as internalized, ranges from 87 to 99%, with a score increasing with size. Specificity, i.e. the part of external targets correctly classified, ranges from 77 to 94%, increasing with size.

**Table 1 :** Estimation of the efficiency of the algorithm for detection and classification of microbeads. Detection rate is the number of detected targets among the targets identified by human eye. Error rate is the number of non-targets objects detected by the algorithm. Sensitivity is the number of targets correctly classified as internalized by the machine among the targets internalized as determined by the observer. Specificity is the number of targets correctly classified as external by the machine among the targets external as determined by the observer.

| Diameter of target | Bead<br>3μm | Bead<br>6μm | Bead<br>10μm | Droplet<br>3μm | Droplet<br>5.5μm | Droplet<br>8μm |
| --- | --- | --- | --- | --- | --- | --- |
| Detection rate | 88% | 93% | 97% | 83% | 92% | 94% |
| Error rate | 0% | 4,7% | 0,6% | 1,2% | 0.5% | 0.3% |
| Sensitivity | 99% | 95% | 92% | 88% | 87% | 87% |
| Specificity | 89% | 94% | 90% | 77% | 89% | 90% |

### Influence of target size on phagocytic efficiency

The database structure allows us extracting complex information from our experiments for a large number of cells (n >3000). More specifically, we used the algorithm to study the influence of the target size on the phagocytosis efficiency. We first measured the phagocytic efficiency of monodisperse populations of droplets (diameters: 3μm, 5.5μm and 8μm), both for the case of uncoated targets and targets coated with fluorescent IgGs. The droplets were saturated with IgG, to prevent the formation of contact clusters of IgG at the surface of droplets. Since the size of the clusters would depend on the size of the droplet, cluster formation would interfere with the assessment of a size effect on phagocytosis. For all experimental conditions, we worked with more targets than the theoretical uptake for a given size. **Figure 5A** demonstrates that the specific, FcγR-mediated, phagocytosis is 5 times more efficient than the non-specific uptake measured for naked droplets. The graph shows that both non-specific and specific internalization is higher for smaller droplets, which is confirmed in **Figure 5B** by the linear increase of the phagocytic index with the inverse of target surface. Moreover, this scaling with the surface of targets prevails for both coated and uncoated targets, which means that the total engulfed surface is independent of the size of targets. Finally, **Figure 5C** shows that the distribution of the total wrapped surface is independent of the size range of monodisperse droplets.

**Figure 5:**
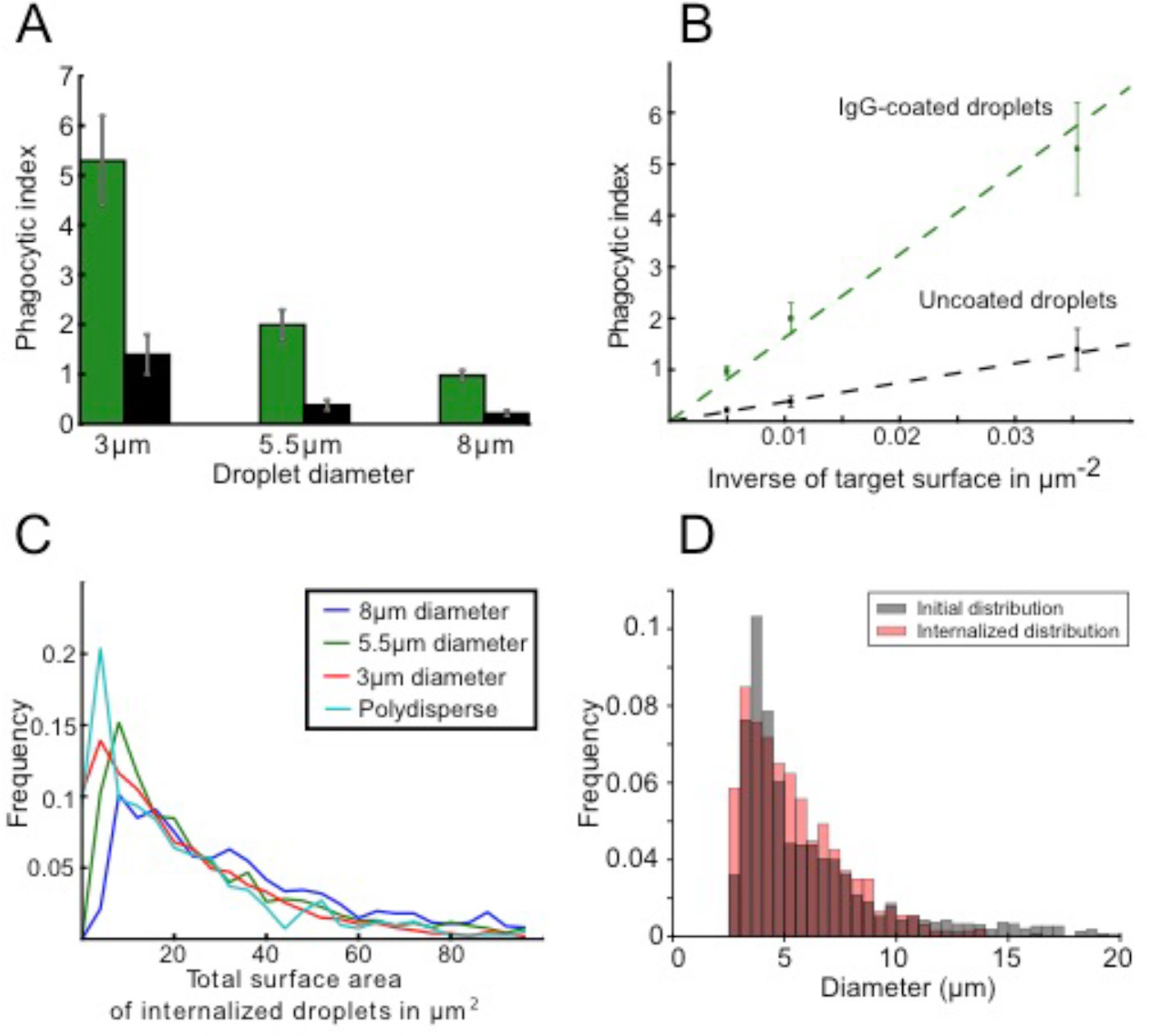
(A) Phagocytic index for 3 sizes of monodisperse droplets with and without IgG-coating. (B) Phagocytic index as a function of the inverse of droplet surface. Both coated and uncoated droplets are internalized proportionally to the inverse of target surface. (C) Histogram of total engulfed surface for 3 monodisperse and 1 polydisperse emulsion, obtained by summing the surface area of the droplets in each cell. (D) Histogram of internalized and initial droplets diameter for polydisperse targets, showing no significant difference.

To check for synergetic effects related to the presence of targets differing by size, we performed the experiments and quantified it for the case of naked and IgG-coated polydisperse droplets ranging from 2 to 15 μm. **Figure 5B** shows that the size distribution of internalized droplets is the same as the initial droplet size distribution, and **Figure 5D** shows that the distribution of total wrapped surface in the case of polydisperse droplets is similar to the distributions measured for monodisperse droplets and detailed above.

## DISCUSSION

In this work, we present the design semi-automatic method based on confocal microscopy, image analysis and the use of a database for the quantitative analysis of phagocytosis assays. The algorithm yields comprehensive data on cells, targets and their respective geometric and fluorescence parameters. It automatically discriminates internalized from external particles, and finally stores the relationships between cells and uptaken particles. Currently, the method is able to analyze samples of up to 3000 cells, and is only limited by the microscopic imaging as this part is performed manually. As compared to manual counting, we gain at least one order of magnitude [28,29] in terms of sample size. In addition, unlike methods published so far, the experimental procedure does not require to stain the targets for the segmentation. This means that the method make it possible to analyze the signal associated to non-specific phagocytosis, i.e. the one measured for negative control experiments.

The automation creates three main challenges: segmentation of the cells, segmentation of the targets, and classification of the targets as internalized or external. The procedure is thus based on the image recording over 3 channels (bright field for the targets, fluorescence of Hoechst for the nuclei and of phalloidin for the cytoplasms). The detected nuclei served as seeds that propagate into the cytoplasm and stop at the borders of the cells. Over the remaining channels, proteins of interest can be labeled, and the intensity and localization of their fluorescence are assessed on the images at the two levels. As an example, we imaged the fluorescent anti-body Alexa488-anti-biotin that coat the droplets and stained the cells for Fcγ-receptors using anti-Cd16/Cd32 antibodies, as can be seen in **Figure S4**.

Correctly segmenting the cell is paramount for correct classification, as the main determinant of the classification is the overlap between cytoplasmic and target areas. Our approach here is similar to [24,25], and allows measuring the morphology of the cells. The stained nuclei are segmented first using thresholding; the cytoplasms are then propagated from the nuclei according to the method developed by Jones *et al* [43]. The use of a confocal microscope conjugated to phagocytic cells added several subtleties to this classical approach, since the presence of internalized targets cause dark areas inside the cytoplasm. Firstly, the detection of cell borders from the cytoplasmic stain has to be performed on the basal level of cells, where the signal from the actin cytoskeleton is higher and more uniform. Secondly, the darker areas in the higher plane can then be exploited to confirm the internalization of targets.

Regarding target detection and classification, two main strategies emerge from the literature. The first one is to classify internal and external targets by staining them differently. The cells and targets are fixed at the end of the phagocytic assay, and a staining step labels external targets only. All targets are detected, either using a pre-fixation stain [20–22] or bright field images [23]. The classification is thus performed before the imaging step, and the detection subsequently on both populations of targets. This protocol, while limiting greatly the risk of errors in classification, is only usable on fixed samples. Moreover, if one needs to relate one cell to its internalized targets, it is still necessary to segment the cell contours and to examine the overlap between the cytoplasms and the targets. We chose instead to use the overlap between the segmented cells and targets as the main criterion for classification, as [24,25]. The difference in our approach was to use the bright field images at the two planes Z_high_ and Z_low_ to segment the targets. It has two advantages: first, to be efficient even in crowded environments, where threshold-based segmentation would fail to separate targets, and second, to exclude out-of-focus targets, reducing false-positive rates, while simultaneously reducing the number of fluorescence channels necessary for the automation. This strategy for cell and target segmentation requires only two stains for cell labeling and none for the targets, when most of the alternative protocols required at least three, either two for the cells and one for the targets [24,25] or one for the cells and two for the targets [20–22]. Moreover, the phagocytosis of targets without staining of any kind can be assessed using this strategy, as it requires no labeling and provides large number of cells to analyze to compensate for the low uptake of unopsonized particles. In our case, experiments were performed of fixed samples, but as long as the dyes are compatible with cells, the same protocol could be used on images taken from live cells.

Classification of targets as internalized or not is performed on segmented cells and targets, using two criteria: the overlap between cytoplasms and targets, and the low cytoplasmic staining at the location of the target. The second criterion requires confocal imaging, and is useful to exclude adherent targets or targets being internalized as the samples were fixed, in which case the cytoskeleton is well developed around the target, or targets above the cell, where the overlap is positive but the cytoplasmic staining is visible inside the target projected area. The sensitivity, i.e. the part of targets correctly classified as internalized, ranges from 92 to 99% for beads and around 87% for droplets, with a score decreasing with size. The specificity, i.e. the part of targets correctly classified as external, ranges from 89 to 94% for beads and from 77% to 90% for droplets. Because IgGs cluster at the contact site with cells, droplets adhere more to macrophages than beads do. Adherent droplets often surround macrophages, increasing the number of false positive and false negative during the classification step as compared to beads. The two measurements depend on one another: when criteria for internalized targets are made more precise, more and more targets are incorrectly classified as external, and the gain in sensitivity is compensated by a loss in specificity. Since we favored sensitivity over specificity, we tend to underestimate the phagocytic indexes. However, the error on the phagocytic index due to misclassification is lower than the variability from one biological sample to another, which diminishes the gain that would be obtained by optimizing further the classification algorithm.

The final database stores measurements for cells, targets, and the relationship between the two. It enables, for any cell to have access to the measurements made on the targets it contains, or conversely for any target to have access to the measurements on the cell it is in, or on the other targets internalized with it. The ability to connect the cells and targets make possible to draw a lot of information from assays using polydisperse targets. Here it was used to compute for each cell the total area of engulfed targets, and confirm that in a population of cells, the distribution of internalized total area does not depend on the area of the targets. It can be used to sort cells based on the properties of the targets they have internalized and thus broaden the scope of hypotheses that can be tested.

### Target size and phagocytosis

Experiments with monodisperse and polydisperse emulsions lead to apparently contradictory results: smaller monodisperse targets are more often internalized, whereas the size distribution is unchanged for polydisperse targets, the cells showing no preference for smaller targets. The phagocytic index dependence in size is the same for coated and uncoated targets, and it is linearly proportional to the inverse of the targets’ area for both, differing only by the slope. Thus, the product of the number of internalized objects and of the area of said objects, i.e. the mean total surface area internalized by cells, is a constant. This constant however depends on the coating of the targets, since non-specific and specific curves have a different slope (**Figure 5B**).

In 1988, Simon *et al*. [28] proposed a simple geometrical model, postulating that membrane surface availability was the limiting factor of phagocytosis. During phagocytosis, the engulfed particle is wrapped closely in the phagosome, and the available cell membrane is consequently reduced, while the cell volume increases. Simon *et al*. measured the initial surface and volume of neutrophils, and assumed that, at maximum uptake, the cells would round up to minimize their surface. They derive a theoretical phagocytic index, depending on measured initial cell volume and surface, on target size and on addition of surface from compartments reservoirs. This model agrees with our observations that the total surface area of internalized objects is constant.

We compiled the results from the literature when quantitative phagocytic index were given or could be computed, and plotted them on **Figure 6**, as a function of the inverse of target surface area, along with the model [28] and our own measurements. While results for targets within 3-10μm ranges are consistent with our own and with each other, results for smaller targets are not. However, the highest measured phagocytic index aligned well with those of larger targets and with the theoretical prediction. At smaller sizes, the theoretical uptakes is above a hundred of targets per cell, yet, cells were not always provided with that many particles. In **Figure 6**, we circled in red every experiment where the cells were not provided with at least five times the theoretical maximum of targets. Circled results are also the ones that differed from theoretical prediction. It is to be noted that **Figure 6** brings together results obtained on 6 types of cells, adherent or in suspension, 4 types of objects, with 4 experimental methods (see **Table S1** for details)

**Figure 6:**
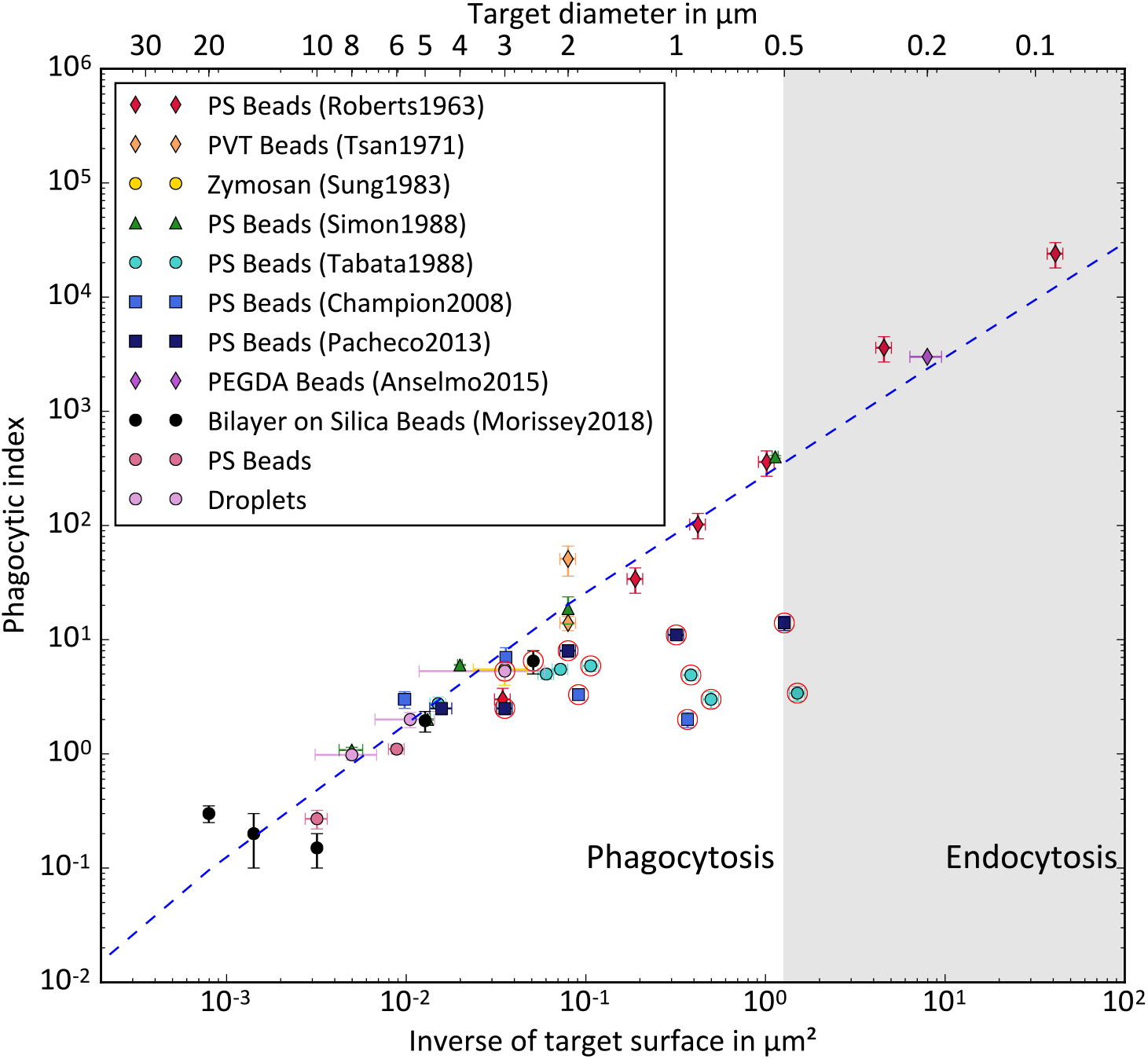
Phagocytic index as a function of the inverse of target surface as collected from the literature [15–19,26–29] and our experiments. Theoretical prediction from Simon *et al*. [28] is shown in dotted blue line. Experiments where the cells were not provided with at least 5 times the theoretical number of targets are circled in red.

Previous experiments addressing the influence of size on phagocytosis were always performed with monodisperse objects, presumably due to the constraints of data collection. The main improvement of the technique we designed is to give access to the individual objects measurements for both cells and targets, as well as their relationship. Thus, we could perform experiments using polydisperse emulsions and subsequently measure the size distribution of internalized objects. Should the cells prefer internalizing targets of a specific range of size, the distribution of internal targets would be skewed to reflect the preference. When presented with a polydisperse emulsion, cells display no preference for one size over another, since the size distribution of targets is conserved by internalization.

We can also compute, at an individual level, the total area of the targets internalized by each cell. For the three sizes of droplets, the distribution of mobilized surface per cell is similar, with more small targets internalized per cell as to compensate for their smaller surface.

Taken together, these results points towards a surface limitation rather than a preference for a target diameter: cells internalize targets they encounter up to their available surface, regardless of the target size as long as the surface can be mobilized. This is also consistent with experimental results from [11], that showed available membrane is a limiting factor for phagocytosis and that reaching the surface limit triggers degranulation. It is interesting to note that this relationship is maintained for uncoated objects internalized by macrophages through other receptors, with a lower total surface limitation: it could point towards membrane availability being dependent on receptor activation, for example through the exocytosis of surface [11,28,44,45]. One of the parameters regulating the selectivity of phagocytosis could then be the controlled delivery of membrane surface by activated receptors.

## CONCLUSION

This method combines the advantages of several of the previous protocols: no staining of the targets is required, geometrical and intensity parameters are measured on cells and targets. It also adds a layer of information by storing the relationship between the objects, allowing much more complex observations. The data collected during experiments on the role of size in phagocytosis shed a new light on results previously reported in the literature. Using polydisperse emulsion droplets and individual target and cell measurements, we concluded that macrophages engulfed targets regardless of size until they reach a fixed internalized surface. A review of over fifty years of experiments on the role of size concurs to show this surface limitation as the main predictor of the phagocytic uptake regardless of cell types or targets’ nature.

## AKNOWLEDGMENTS

We thank Nicolas Carpi (Bio6 team, UMR 144, Institut Curie) for his help on the software. This work has received support of “Institut Pierre-Gilles de Gennes” (Laboratoire d’excellence : ANR-10-LABX-31, “Investissements d’avenir” : ANR-10-IDEX-0001-02 PSL and Equipement d’excellence : ANR-10-EQPX-34). The authors acknowledge funding from the ANR JCJC (Phagodrop). We thank XX and XX for fruitful discussions about the work, and their help at improving the readability of the manuscript.

## SUPPLEMENTARY INFORMATION

### 1. Size distribution of the oil droplet samples

**Figure S1:**
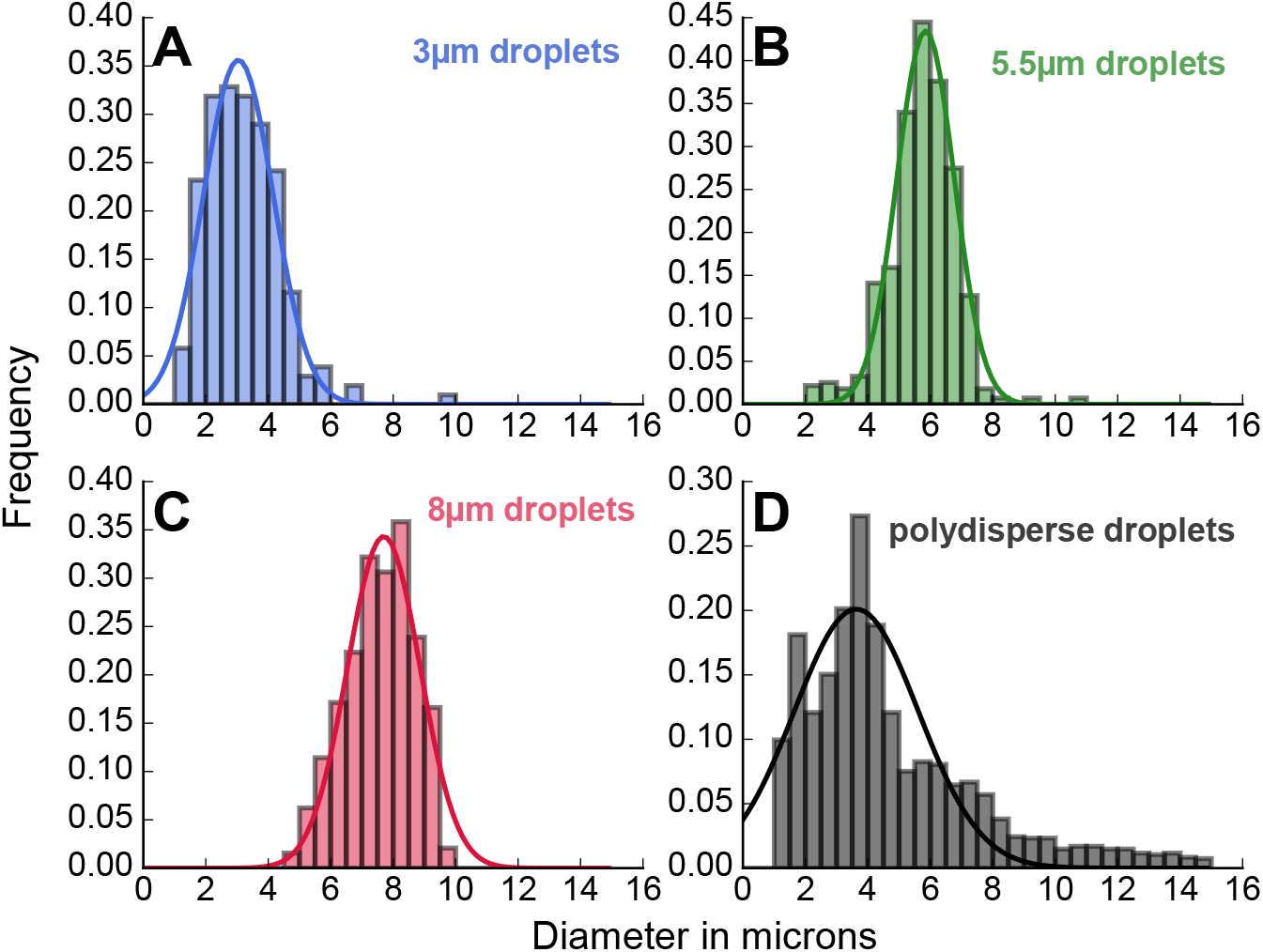
Histogram and Gaussian fit for (A) 3μm diameter droplets, (B) 5.5μm droplets, (C) 8μm droplets, (D) polydisperse droplets.

### 2. Brightfield imaging of droplets and beads with the confocal microscope

**Figure S2:**
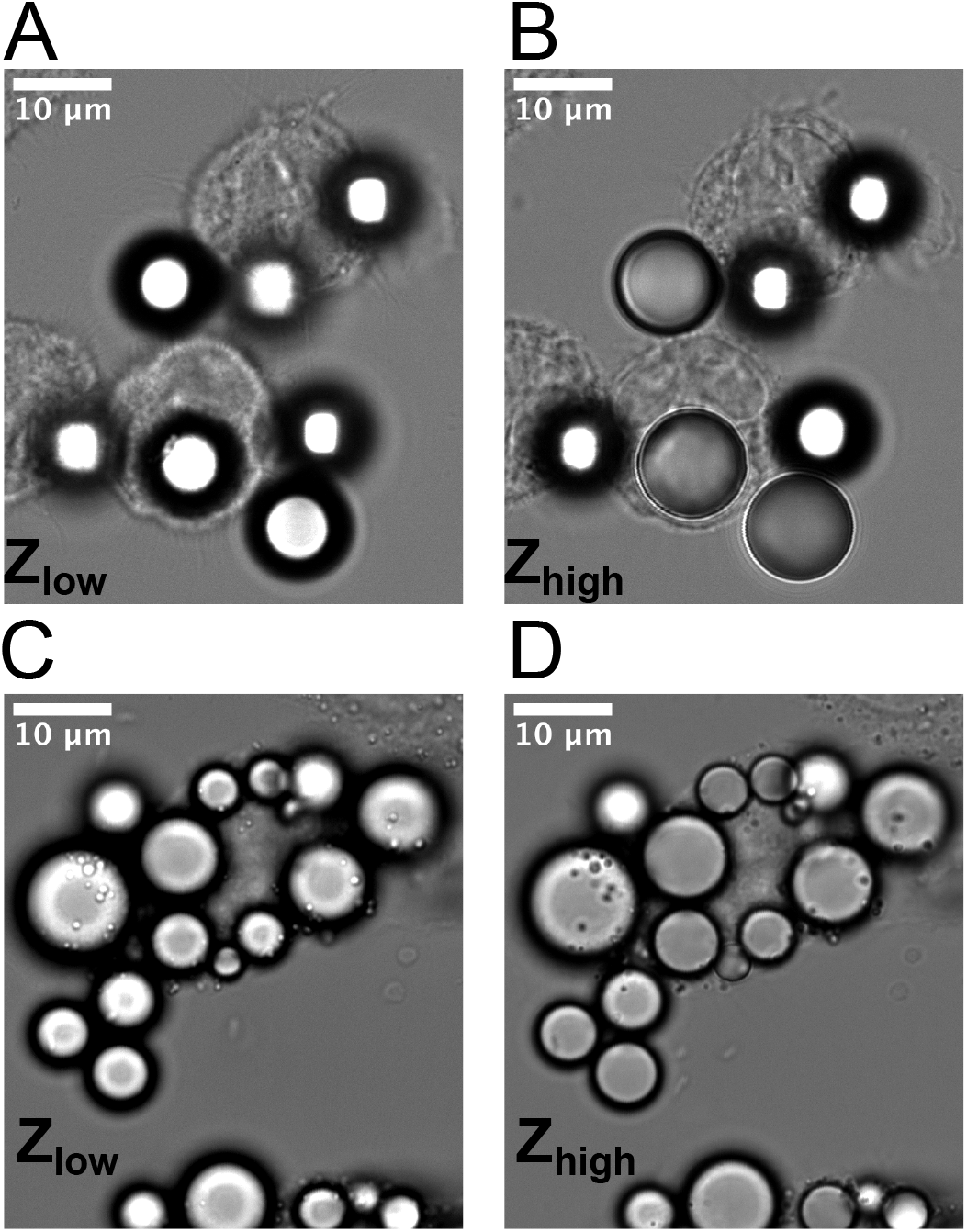
Bright field images of (A) Polystyrene beads at Z_low_, (B) Polystyrene beads at Z_high_=5μm, (C) Oil droplets at Z_low_, (D) Oil droplets at Z_high_=4μm

### 3. Detailed description of the database

The database generated by CellProfiler contains the measurement data for four types of objects, grouped in tables Per Object and Per Image, and the relationship tables connecting the objects to each other. Object types are nuclei, cytoplasms, cytoplasms minus targets and targets.

A unique Image Number identifies each field of view, and is the primary key of the Per Image tables. The Image Number and Object Number form the primary key of objects.

Each child nucleus is related to exactly one parent cytoplasm, each parent cytoplasm to exactly one child cytoplasm minus targets. Each parent cytoplasm can be related to one or more children internalized targets. The Per Object tables contain the intensity and shape measurement for each object, as well as the identifier of its parent objects or the number of its children objects.

**Figure S3:**
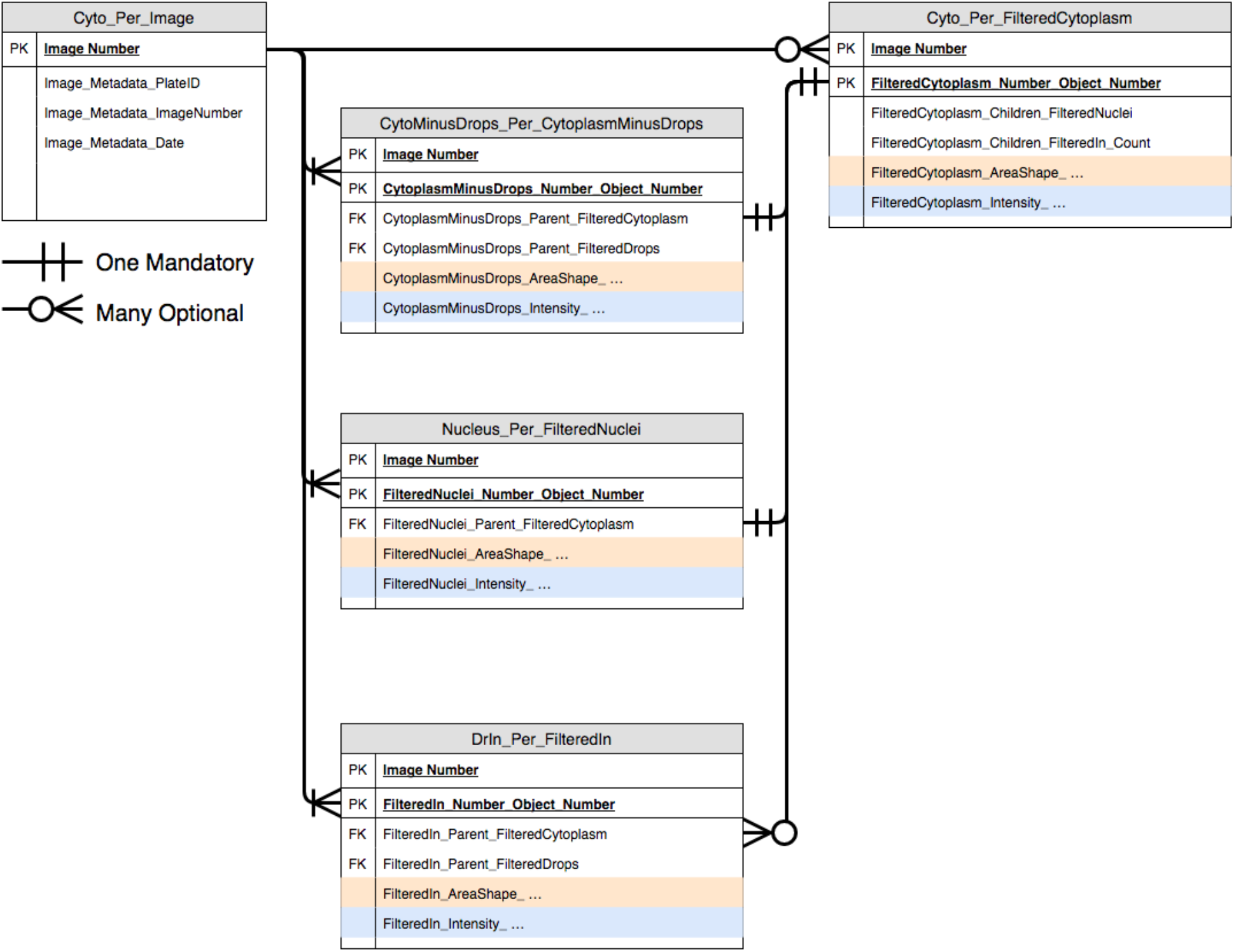
Functional diagram of the database structure. The AreaShape and Intensity fields, not represented fully here, contain 18 geometric measurements and 15 intensity measurements per channel, respectively.

### 4. Example image with IgG and FcγR staining

Fixed cells are incubated for 20 minutes in NH4Cl 50mM, permeabilized with 0.1% Triton X-100 for 10 minutes, saturated with 4% Bovine Serum Albumin for 30 minutes. Fcγ Receptors were stained using Purified Rat Anti-Mouse CD16/CD32 (BD Pharmingen Bioscience, ref 553142) 1μg/chamber for 15 minutes. After washing with 4%BSA (Sigma Aldrich, ref A3059-10G), cells were stained with secondary antibody Goat anti-Rat Alexa 647 8μg/ml (Life Technologies, ref A-21247), Hoechst 33258 4μg/ml (Life Technologies, ref H H1399) and Atto 550-phalloidin 20μM (Sigma Aldrich, ref 19083) for 30 minutes.

**Figure S4:**
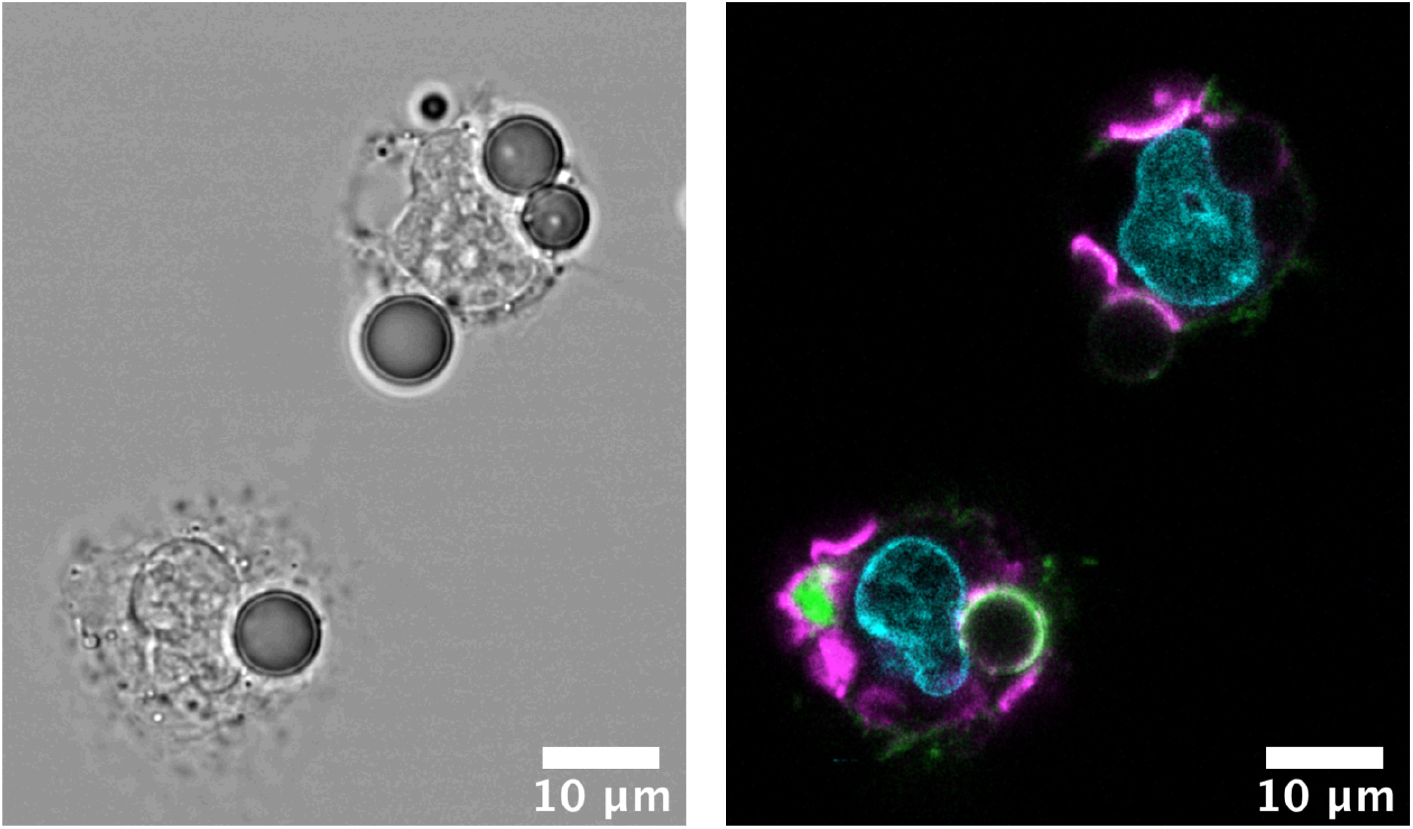
Brightfield and fluorescence confocal image of Alexa488-anti-biotin coated droplets (green) and macrophages stained with anti-Cd16/32 for Fcγ Receptors (magenta). Patches of receptors can be observed at the contact zone with droplets that are visible on the image, or with droplets that were detached during the rinsing steps.

### 5. Details of the meta-analysis on the influence of size on phagocytosis

**Table S1 :** Table of the main parameters of the phagocytic assays for the data plotted in **Figure 6**.

|  | Reference | Cell Type | Target type | Presented targets to theoretical uptake ratio | Method |  |
| --- | --- | --- | --- | --- | --- | --- |
| 1 | [15] | Leucocytes<br>(84% neutrophils) | PS beads | Unspecified | Spectrophotometry | suspended |
| 2 | [16] | Neutrophils<br>Alveolar<br>Macrophages | PVT Beads | 10 - 80 | Spectrophotometry | adherent |
| 3 | [26] | Peritoneal<br>Macrophages | Zymosan | 10 | Microscopy | adherent |
| 4 | [28] | Neutrophils | PS Beads | 5-10 | Microsection+TEM | suspended |
| 5 | [27] | Peritoneal<br>Macrophages | PS Beads | 0.002-10 | Microscopy | adherent |
| 6 | [18] | Rat alveolar<br>Macrophages | PS Beads | Unspecified | Cytometry | adherent |
| 7 | [19] | RAW264.7 | PS Beads | 0.1-10 | Cytometry | adherent |
| 8 | [17] | J774 | PEGDA<br>nanoparticles | 100 | Spectrophotometry | adherent |
| 9 | [29] | J774 | Lipid Bilayer<br>on Silica<br>Beads | 2-200 | Microscopy | adherent |

